# Cellpose-SAM: superhuman generalization for cellular segmentation

**DOI:** 10.1101/2025.04.28.651001

**Authors:** Marius Pachitariu, Michael Rariden, Carsen Stringer

## Abstract

Modern algorithms for biological segmentation can match inter-human agreement in annotation quality. This however is not a performance bound: a hypothetical human-consensus segmentation could reduce error rates in half. To obtain a model that generalizes better we adapted the pretrained transformer backbone of a foundation model (SAM) to the Cellpose framework. The resulting Cellpose-SAM model substantially outperforms inter-human agreement and approaches the human-consensus bound. We increase generalization performance further by making the model robust to channel shuffling, cell size, shot noise, downsampling, isotropic and anisotropic blur. The new model can be readily adopted into the Cellpose ecosystem which includes finetuning, human-in-the-loop training, image restoration and 3D segmentation approaches. These properties establish Cellpose-SAM as a foundation model for biological segmentation.

## Introduction

The most important aspect of biological software is that it works well in the hands of biologists. This typically requires good performance on new data, acquired in new experiments, possibly in new tissues or using new stains or new microscopes. Such data is often outside of the input distribution that models have been trained on. Algorithm developers can make an effort to develop models that anticipate user needs, but it is difficult to predict what new datasets will require segmentation. Instead, developers may specifically focus on methods that can be proved to generalize well out-of-distribution. Pursuing this goal is not straightforward, because many of the existing datasets and challenges contain homogeneous datasets, in which test images are very similar to train images [1, 2]. Successful models on these datasets are those that can best memorize training patterns and convert that knowledge into segmentations.

Converting knowledge into segmentations is not straightforward. Previous versions of Cellpose excel at this [3–5]. As we show below, they even outperform the latest foundation models such as the Segment Anything Model (SAM), that has been recently adapted to biological segmentation by multiple groups [6–9]. Models may perform better based on their architecture, loss function and post-processing steps [10]. For example, the Cellpose loss function and post-processing has often proved advantageous, outperforming models like Stardist and Mask R-CNN when trained on biological data [3, 11, 12]. Other frameworks such as prompt-based segmentation have recently been developed in computer vision and adapted to biology [6–9], but it is unclear if these frameworks perform as well as Cellpose, due to difficulty interpreting benchmarks in their respective studies and because the training strategies and model backbones (U-Net, Transformer etc [10]) were also varied. In this study, we show that replacing computer vision segmentation frameworks with the Cellpose framework gives a major boost in performance in foundation models such as SAM.

Foundation models like SAM do have some unique properties. Due to being pretrained on very large datasets, these models develop representations that have strong inductive biases. Such biases can be highly beneficial for out-of-distribution generalization, especially when finetuning on limited data [13, 14]. In a “best case” scenario, foundation models may “understand” what tasks they are asked to do, and use their general purpose computations to complete these tasks, not unlike how a human may approach a new task. Metaphors aside, such models still require mechanisms and computation to transform knowledge into segmentations, a task that Cellpose is uniquely well-suited to achieve.

Thus, we designed a new Cellpose-SAM model that combines the Cellpose framework with the pretrained SAM weights, and we make it available locally^1^ and online^2^. Below, we start by describing the model design. Then we explain why inter-annotator agreement is not a true upper bound for model performance, and show that a hypothetical “average” or “consensus” annotation has about half the error rate. We show that Cellpose-SAM approaches this hypothetical bound for the Cellpose test set, while previous models do not exceed inter-annotator agreement. From there we describe all the augmentations that were used for Cellpose-SAM; while these do not necessarily improve performance, they allow for much more flexibility in using Cellpose-SAM across a range of datasets. Then we show that Cellpose-SAM can be adapted more quickly to new datasets, which also holds in 3D. Finally we show that Cellpose-SAM can be retrained as a panoptic segmentation model for joint segmentation and semantic classification.

## Results

### Model design

Our goal in Cellpose-SAM was to take advantage of SAM as a foundation model that was pretrained on a large dataset. This pretraining allows SAM to learn the structure of natural images and this knowledge can be beneficial when training the model on a new task. However, SAM also has design choices that make it poorly suited for dense image segmentation. We wanted to replace those weak points with the advantages provided by the Cellpose framework. Briefly in Cellpose, a U-net type neural network is used to predict a set of vector flows, which form an intermediate representation of the segmentation [3] (Figure 1a). These vector flows can be iterated in parallel at every pixel by gradient tracking to produce a set of masks. Conversely, the flow fields can be constructed from the masks as training data for the neural network. By contrast, SAM predicts masks in an image sequentially one-by-one, based on prompts given to the algorithm either in the form of point(s), a box, or text and which get further processed by specialized modules [6] (Figure 1b). To densely predict all the cells in an image, biologically-adapted versions of SAM add another set of modules and neural networks [7–9] (not shown in Figure 1b). This strategy for generating dense segmentations is cumbersome and requires careful tuning of many different parts.

**Figure 1:**
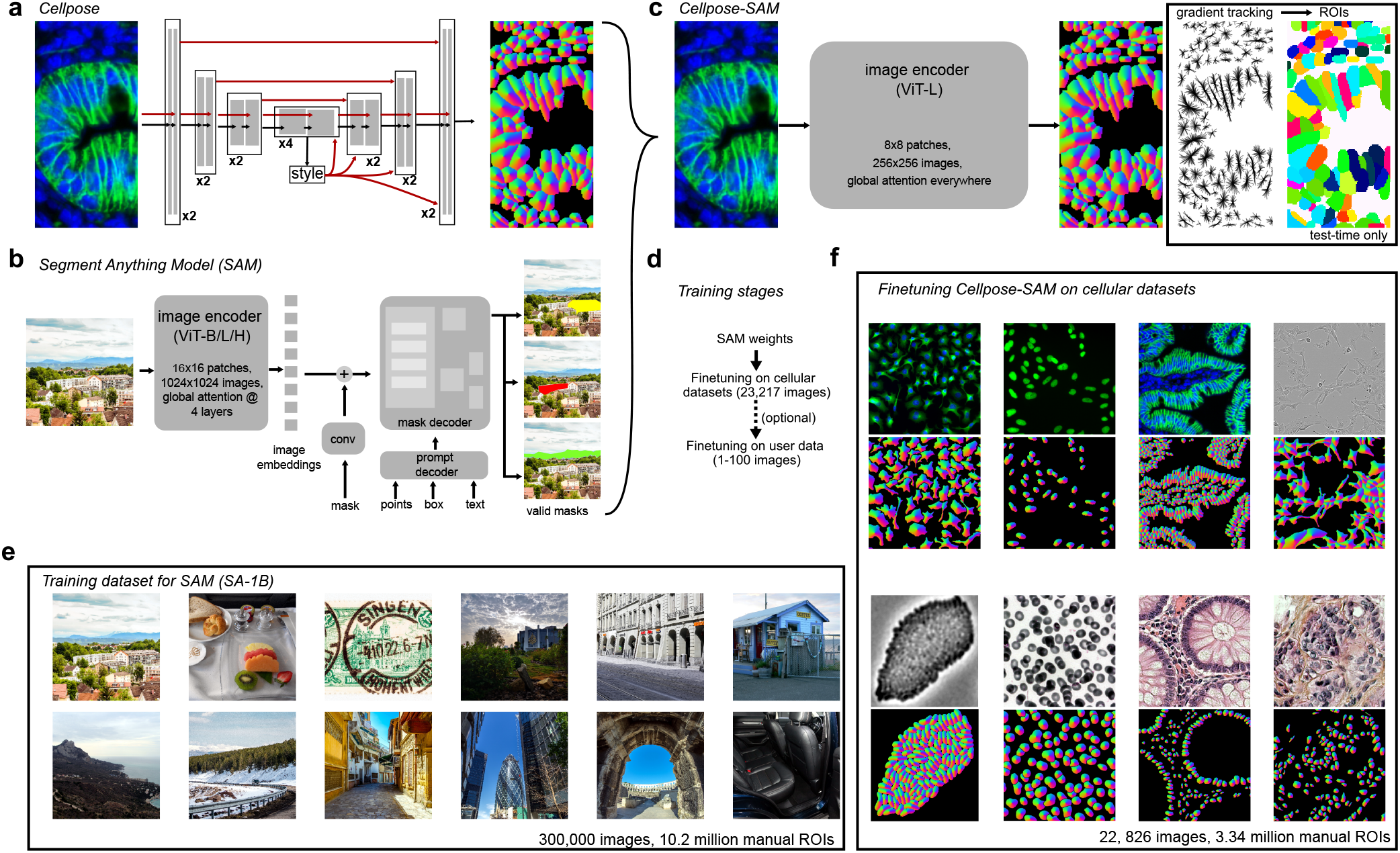
Defining and training the Cellpose-SAM model. **a**, Schematic of the U-net backbone used in the original Cellpose model with skip connections and global style vector. Shown on the right is the representation of the flow vectors that Cellpose predicts as an intermediate to mask reconstruction. **b**, Schematic of the Segment Anything Model that includes an encoder based on a ViT backbone with a few modifications, as well as complex mask and prompt decoders required for both training and inference. **c**, Schematic of the Cellpose-SAM model combining a customized encoder backbone based on SAM, with the flow field prediction and gradient tracking of Cellpose. **d**, Training stages for Cellpose-SAM. **e**, Example images used to train the original SAM model. **f**, Example images and flow fields for the updated training dataset for Cellpose-SAM.

We decided to instead eliminate entirely the decoder modules of SAM and use the image encoder exclusively, which contains a majority of the parameters (305M out of 312M, Figure 1c). From the encoder output, we directly predicted the vector flow fields of Cellpose, without any intermediate modules (Figure 1c). In addition, we made a few modifications to the encoder itself. The default 1024×1024 image inputs and 16×16 patch sizes of SAM were designed for high-resolution photographs [6, 15]. We reduced this to 256×256 and 8×8, which required us to adapt the position embeddings and patch embedding filters via appropriate subsampling. These modifications improved runtime performance and ensured that more computation is dedicated to each region of the image. We also reverted the local attention layers of the custom ViT transformer from SAM back to the default global attention of ViT-L, with almost no runtime penalty [16].

Despite making these modifications, we were still able to initialize Cellpose-SAM with the SAM weights pretrained on the SA-1B dataset [6] (Figure 1de). The model was then trained on an updated dataset of cells and nuclei containing 22,826 train images with a combined 3,341,254 training ROIs. This dataset combines major currently available datasets: Cellpose, Cellpose Nuclei, Omnipose, TissueNet, LiveCell, YeaZ, DeepBacs, Neurips 2022, MoNuSeg, MoNuSAC, CryoNuSeg, NuInsSeg, BCCD, CPM 15+17, TNBC, LynSec, IHC TMA, CoNIC, PanNuke [1–3, 17–36] (Figure 1df, Figure S1). We continued to use the Cellpose3 mixing probabilities for down-weighting homogeneous datasets with many images [5].

### Model validation

Since we are especially interested in generalization performance, we wanted to choose a test dataset in which images are relatively different from those in the training set. To determine this, we extracted feature vectors that describe the styles of images [5, 10, 37], and correlated them between pairs of images (Figure 2a). In most datasets, the style vectors were highly-correlated between all pairs of train/test images (Figure 2b), matching our subjective experience of these images. The Cellpose dataset alone contained a high amount of variability between train/test exemplars, leading to low correlations between the styles of train/test images. We therefore chose to focus our benchmarking efforts on the Cellpose test set.

**Figure 2:**
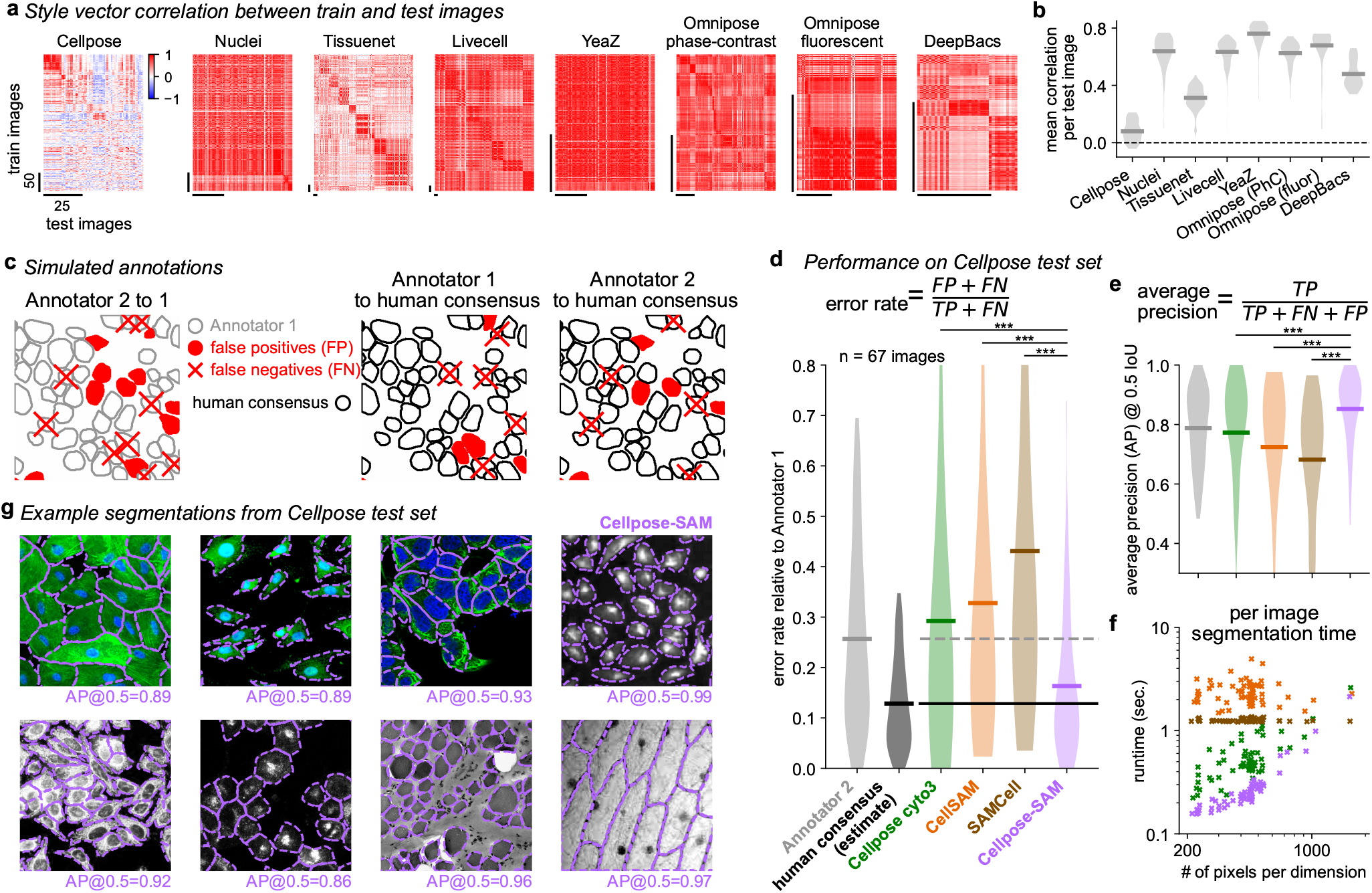
Performance relative to other methods and to humans. **a**, Simulated annotations to illustrate human-to-human variability and the human consensus as a hypothetical absolute bound on performance. **b**, Performance on the updated Cellpose test set shown as error rates (left) and average precision (right). Per-image segmentation time on an A100 GPU (bottom). **c**, Example segmentations of Cellpose-SAM, taken from the updated Cellpose test set.

Before we evaluate performance, we must set our expectations. In general computer vision applications, improvements on the order of a few percent can be considered major steps [38–40], and comparisons to human performance can help in setting targets for improvement [41]. Previous biological segmentation models, including Cellpose, have described performance as matching inter-annotator agreement [4, 7, 42], which is defined as the “performance” of one annotator when benchmarked against another (Figure 2c). While this can be a useful reference, it is not a true upper bound, because both annotators make mistakes. Careful consideration shows that if we model each annotator as making errors randomly starting from the same underlying ground truth or “human consensus”, then the number of errors for inter-annotator comparisons is less than twice as large as that between each annotator and the human consensus, and approaches the two-fold limit when the errors are relatively small (Figure 2c, see Methods for exact calculations). Thus, setting the human consensus estimate for errors at half the inter-annotator rate results in a performance bound on performance that automated models may be able to reach.

To determine the inter-annotator variability we relabeled the Cellpose test set using a different annotator. Relative to the original Annotator1, the error rate of Annotator2 was 0.257 (Figure 2d). Halving this error rate gives the human consensus estimate at 0.128. Previous models, like Cellpose3 and CellSAM approach the inter-annotator error rates, at 0.292 and 0.328 respectively (Figure 2d). Cellpose-SAM however, achieves error rates of 0.163, substantially below inter-annotator agreement, and approaching the human consensus estimate. Note this was possible despite the models being trained on data from Annotator1 exclusively. Rather than being “fooled” by the occasional errors of Annotator1, Cellpose-SAM reverts to its inductive biases to selectively learn the generalizable structure in the data. We can draw similar conclusions using the average precision @ 0.5 intersection-over-union (AP @ 0.5 IoU) score (Figure 2e). While this metric is more widely used than the error rate, it does not scale linearly with the number of errors, so we cannot easily estimate the human consensus bound. Finally, we measured runtime performance (Figure 2f, Table S1). Despite having 50x more parameters than Cellpose, Cellpose-SAM is the fastest of the models considered here when benchmarked on a per-image basis. This is due to a few factors: 1) modern GPUs have specialized tensor cores that can vastly accelerate transformer computations; 2) the post-processing times account for a substantial fraction of the runtime; 3) Cellpose-SAM does not need to make two passes for every image like Cellpose 1/2/3, which need to first estimate the sizes of cells using a size model. As we describe below, Cellpose-SAM runs natively on images at a wide range of resolutions.

We also evaluated the performance of Cellpose-SAM on other public datasets that we used for training. All of these datasets are much more homogeneous than Cellpose, thus providing more images of the same type for training, and a much higher chance that images in the test set are similar to those in the training set (Figure 2a, Figure S1). Thus, models are evaluated mainly for their in-sample generalization, and even models trained from scratch (like previous versions of Cellpose) can perform well given enough training data. Cellpose-SAM outperformed or matched other models on all datasets, but the performance gaps were typically smaller than those we reported above on the Cellpose test set Figure S2. We conclude that Cellpose-SAM especially shines on out-of-sample generalization, which is likely also the crucial property needed by users applying it to their own data.

### Invariance and robustness

To further drive our goal of increasing generalization, we were able to make the model robust to common image manipulations without loss of performance, thus simplifying the user experience. For example, users of Cellpose previously had to indicate which channel of an image represents the nucleus, and which channel represents a cytoplasmic or membrane marker. While this is not in itself an onerous task, it can lead to some confusion, especially when benchmarking against Cellpose, with some public challenges explicitly designed to break the order of image channels by random channel permutations [22]. Thus, we trained Cellpose-SAM to be entirely channel order invariant, by randomly permuting channels at train time (Figure 3a). Note this does effectively withhold information from the model: specifically information about which channel represents which stain. Nonetheless, Cellpose-SAM had no performance loss from this modification (Figure 3a). Similarly, users of Cellpose previously had to indicate the average cell diameters in an image, or rely on a built-in size estimation method. This also led to confusion in benchmarks by other studies [22], and we therefore trained Cellpose-SAM at a range of image diameters ranging from 7.5 pixels to 120 pixels. At test time, Cellpose-SAM runs natively on the provided images without resizing, and maintains its high-performance, with a small performance loss at small cell diameters, where information is effectively lost by downsampling (Figure 3b).

**Figure 3:**
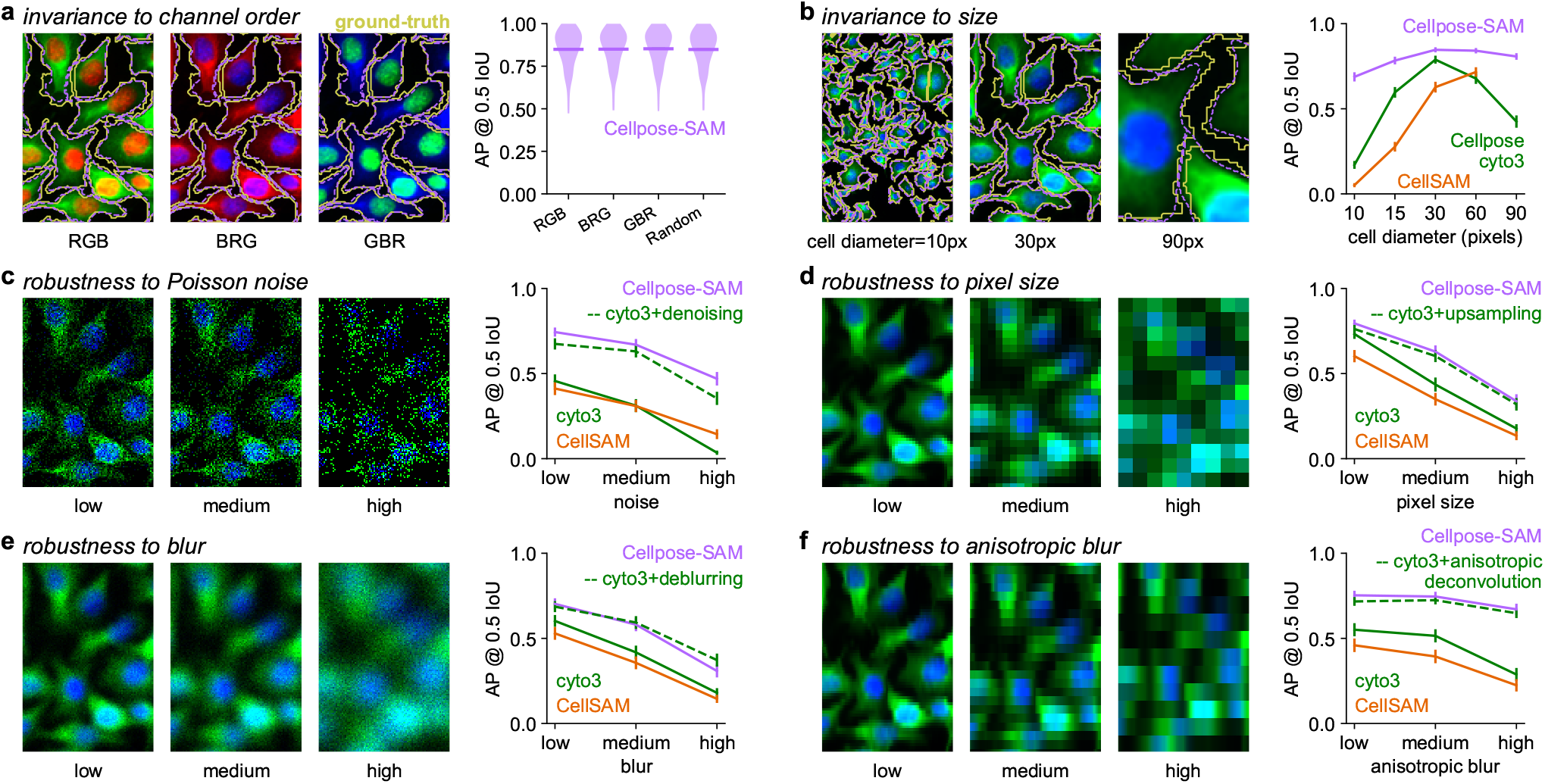
Quality of life improvements for Cellpose-SAM. **a**, Cellpose-SAM is invariant to channel order, because it was trained with channel shuffling. **b**, Cellpose-SAM is mostly invariant to image resizing because it was trained with a wide range of image augmentations. Note that previous Cellpose versions relied on an intermediate size prediction model. **c-f**, Cellpose-SAM is robust to **c** Poisson noise, **d** blurring, **e** image downsampling and **f** anisotropic blur, because it was trained on such images. Note that the previous version of Cellpose relied on intermediate image restoration models to achieve similar performance.

To further take advantage of the high capacity of Cellpose-SAM, we trained it on a set of image degradations that are common in microscopy, and which we previously used in the Cellpose3 study. Specifically these degradations are the addition of per-pixel noise (“shot noise”), downsampling or increasing pixel size and (an)isotropic blurring (Figure 3c-f). Previously we had found that an additional image restoration step was required on such images for best performance [5]; directly predicting the segmentation from the noisy image was inferior for the U-Net based version of Cellpose. The additional restoration step was however not required for Cellpose-SAM. Across all types of image alteration, the Cellpose-SAM model performed as well as our best previous image restoration models (Figure 3c-f). Note that these restoration models are individually trained for each type of image degradation, while a single Cellpose-SAM model was used for all the benchmarks in this entire study. Thus, Cellpose-SAM can run out-of-the-box on images that have been acquired with varying levels of image degradation, at different pixel sizes or in arbitrary channel order, substantially simplifying the logistics typically associated with setting up an image segmentation pipeline.

### Finetuning Cellpose-SAM in 2D and 3D and for other tasks

Next we wanted to test the integration of Cellpose-SAM with other aspects of the Cellpose framework, such as the finetuning and human-in-the-loop capabilities. Compared to inference, training and finetuning a neural network can require considerable more hardware resources. For example, other methods like microSAM have quoted computational concerns as preventing a human-in-the-loop approach for their SAM-based model [8]. We do not find similar concerns when testing Cellpose-SAM, due to our customizations enabling the model to run on low resolution images. We find that it is always possible to train Cellpose-SAM with a batch size of one, even on GPUs with relatively low VRAM (8-12 GB) Table S2. To test finetuning performance, we analyze a series of datasets that we did not use for training, specifically BlastoSPIM [43] and PlantSeg [44]. These datasets contain 3D annotations, from which we can extract both 2D ground-truth, and 3D ground-truth. From the PlantSeg categories, we used the “lateral root” and “ovules” datasets which contained higher-quality segmentations. Across datasets, Cellpose-SAM provided a better starting point for finetuning than the cyto3 model and finetuned equally fast or faster, both in 2D (Figure 4a-c) and in 3D (Figure 4d-f). The performance gap generally narrowed with more training data (Figure 4acdf) but in some cases persisted (Figure 4be). The narrowing gap again illustrates the specific strengths of Cellpose-SAM: the model is especially good at generalizing with zero-shot or limited data. When enough labeled data is available for training, good inductive biases are no longer necessary, and models with less generalization capacity can still perform well.

**Figure 4:**
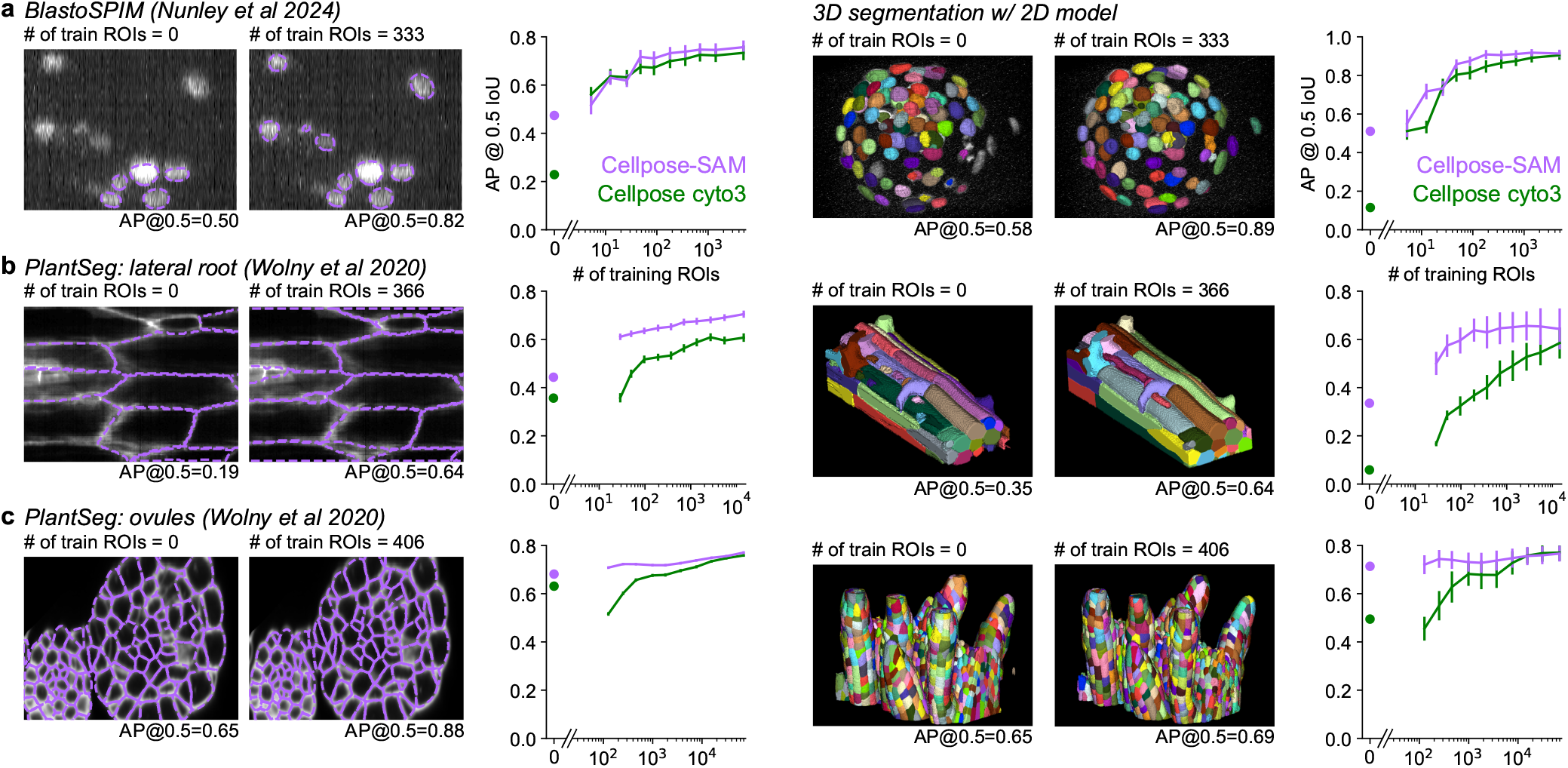
Finetuning performance. **a**, Cellpose-SAM segmentations on the BlastoSPIM dataset [43] either (left) out-of-the-box or (center) after finetuning with a medium number of manually-annotated ROIs. (right) Performance as a function of the number of training ROIs for Cellpose-SAM and Cellpose3. **b-c**, Same as **a** for the “lateral root” and “ovules” categories of the PlantSeg dataset [44]. **d-f**, 3D segmentations extended from 2D predictions using either Cellpose-SAM or Cellpose3. The 3D extension was made using the flow averaging method on XY, XZ and YZ slices from [3]. Same datasets were used as in **a**-**c**.

Generalist models like Cellpose-SAM are often good initializations for other image-based tasks. To demonstrate this, we chose the MoNuSAC 2020 challenge [24], where participants were asked to segment cells in histopathology images and classify them into four classes: epithelial, lymphocyte, macrophage and neutrophil (Figure 5a). This task is often referred to as “panoptic segmentation”. After training, a Cellpose-SAM-based model outperformed the winners of the challenge both before and after the deadline, on nearly all types of cells, and substantially increased the classification accuracy overall (Figure 5b). This example shows what generalist or “foundational” models can achieve when simply trained on new tasks with minimal additional effort required.

**Figure 5:**
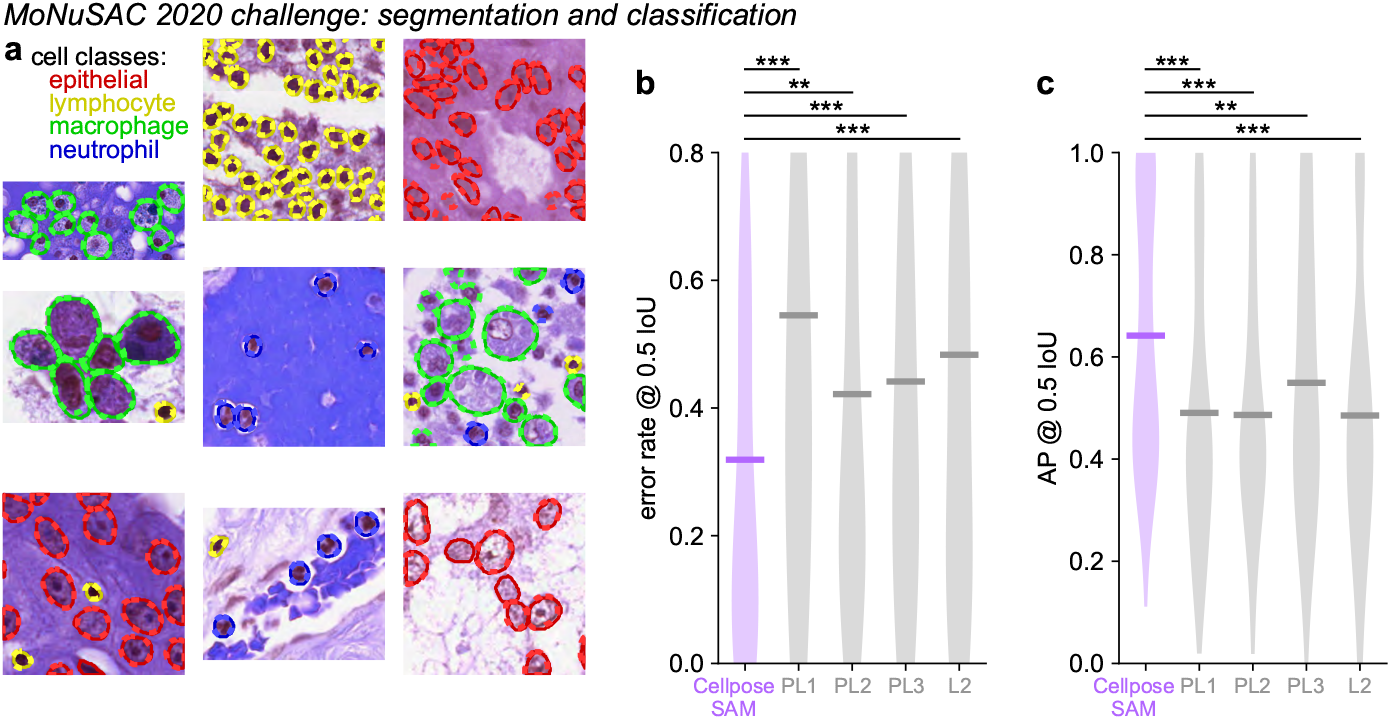
Finetuning Cellpose-SAM for panoptic segmentation. **a**, Representative examples from the MoNuSac 2020 challenge containing four different cell classes. **b**, Segmentation error rate averaged across classes for Cellpose-SAM and four leading algorithms from the challenge (n=85 test images, Wilcoxon signed-rank test). **c**, Same as **b** for average precision.

## Discussion

Here we have shown that Cellpose-SAM can generalize out-of-distribution to a wide range of test images, obtaining performance that is close to the theoretical bound. This was achieved by combining a pretrained foundation model (SAM), which has strong inductive biases due to the broad knowledge built-in to its weights [6], with the Cellpose framework, which is especially well-suited for converting knowledge into segmentations [3]. To demonstrate out-of-distribution generalization, we used the Cellpose test dataset and multiple annotators. This dataset uniquely has a high dissimilarity between train and test images, due to being constructed as a generalist dataset. As such, performance on the test set indicates out-of-distribution generalization. This property, we believe, is the main indicator of how well a segmentation method will work in the hands of its end-users.

Looking ahead, Cellpose-SAM can be used as a basis for many different biological applications. We have shown here that the model can be easily finetuned, and that it can be extended to 3D segmentation. Similarly, 3D segmentations can be extended to 4D (including time) using other methods [45–47]. The model can also be used on images with more than three channels, such as those arising from in-situ sequencing experiments, from “cell painting” or other multi-stain or multi-antibody methods [48, 49], by replacing its three-channel inputs and retraining. The outputs can also be repurposed, for example for classification, as we have shown here [24]. More generally, any task that used previous versions of Cellpose can now use Cellpose-SAM for a boost in performance [50–53]. We especially expect the boost to be high for images that are obtained with non-standard and novel approaches, and for which large annotated datasets are not available for finetuning. Biologists developing new imaging methods or novel molecular approaches may thus especially benefit from the generalization performance of Cellpose-SAM.

## Acknowledgments

This research was funded by the Howard Hughes Medical Institute at the Janelia Research Campus. We thank the authors of [1–3, 17–36, 43, 44] for sharing their datasets.

## Data availability

The ‘cyto2’ dataset is publicly available at https://www.cellpose.org/dataset, and the other datasets were generated and shared by other labs [1, 2, 17– 36, 43, 44].

## Code availability

Cellpose-SAM was used to perform all analyses in the paper. The code and GUI are available at https://www.github.com/mouseland/cellpose. An online version of the algorithm is running on Hugging Face https://huggingface.co/spaces/mouseland/cellpose. Scripts for recreating the analyses in the figures will be available at https://github.com/MouseLand/cellpose/tree/main/paper/cpsam.

## Methods

The Cellpose code library is implemented in Python 3 [54], using pytorch, numpy, scipy, opencv, imagecodecs, tifffile, fastremap, and tqdm [55–62]. The graphical user interface additionally uses PyQt, pyqtgraph, and superqt [63–65]. The figures were made using matplotlib and jupyter-notebook [66, 67].

### Cellpose-SAM network

#### Model architecture

We used a customized version of the ViT-L transformer from the Segment Anything Model [6], which has 24 blocks and an embedding dimension of 1024. Each block contains the standard attention and MLP layers, common to most transformers. We used an input size 256×256 instead of 1024×1024, and we reduced the image patch size from 16×16 to 8×8. To this we add simple input and output operations to convert from pixel space to patch space and back again. Since the SAM model already had strided input convolutions, we adapt these from 16×16 to our 8×8 patch size by downsampling. The position embeddings from SAM were also downsampled by a factor of 2. Next, we changed the attention to global on all layers, rather than only global in layers 6, 12, 18, and 24 like the original SAM. Then, we added a transposed convolution layer to revert the patchified embeddings back into pixel space to predict three pixel maps corresponding to cell probabilities, horizontal and vertical flows (3×256×256, [3]). We used the weights of SAM pretrained on segmentations from photographs. Note that only images not containing humans are shown in Figure 1e. About half of the images in SA-1B do contain blurred humans with blurred faces.

#### Whole-dataset training and evaluation

We trained Cellpose-SAM on the combined dataset of 22,826 images with a combined 3,341,254 ROI annotations (see below for details). All training was performed with the AdamW optimizer [68] using a learning rate of 5e-5, which we empirically found to lead to fast reductions in the training loss. We trained with a batch size of 256, divided across eight H200 GPUs. The learning rate increased linearly from 0 to its maximum value over the first 10 epochs, then decreased by a factor of 10 for the last 100 epochs and another factor of 10 for the last 50 epochs, similar to the SAM training recipe [6]. Epochs were defined as random samples of 800 images per GPU drawn with the mixing probabilities described below, for a total of 6,400 images per “epoch”. The network was trained for 2,000 epochs, which took around 20 hours. The weight decay parameter was set to 0.1, and additional regularization was performed by randomly dropping layers of the image encoder with a 0.4 layer drop rate as previously used for SAM [6].

The loss function was the Cellpose segmentation loss [3]: the mean squared error between the XY flows from the ground-truth segmentation and the predicted XY flows, scaled by a factor of five, added to the binary cross-entropy between the ground-truth cell probability and the the predicted cell probability. The XY flows were computed from the ground-truth masks as described in [5]. During training, all images were normalized such that 0 was set to the first percentile of the image intensity and 1 was the 99th percentile.

In each batch, images were randomly rotated, flipped, and resized with a scale factor logarithmically distributed between 0.25 and 4 relative to a mean cell diameter of 30 pixels, and randomly cropped to an image size of 256×256. Images were randomly converted to grayscale 10% of the time. For single-channel images, grayscale conversion was done by replicating the non-zero channel across all three channels; for H&E images, grayscale conversion was done by taking the mean across all channels and replicating that across all channels; while for images with nuclei, we discarded the nucleus channel and replicated the primary channel. We inverted the image contrast 25% of the time (*x* →1 −*x*). After this operation, the third channel was randomly dropped on 10% of the images, and then the channels of each image were randomly permuted. Finally, the brightness level of each channel (the pixel mean) was randomly perturbed by a random normal jitter of standard deviation 0.2, and the contrast level (standard deviation) was uniformly rescaled by a randomly-drawn factor between -2 and 2. All 256 images in a batch underwent these augmentations.

On 50% of the images in a batch we added four types of degradations: Poisson noise, Gaussian blurring, downsampling, and anisotropic blurring with downsampling. Out of 256 images per batch, exactly 32 had each type of degradation, and the rest of the images were clean (except for the augmentations described in the previous paragraph). The degradation parameters for each of the four conditions were the same as in the Cellpose3 paper [5]. After adding the image degradation, we renormalized the images to again set 0 to the first percentile and 1 to the 99th percentile of the image intensity. In this case, the brightness and contrast augmentations described above were applied after the degradations.

During testing, the tile size was 256. There was no diameter estimation and resizing performed like in previous versions of Cellpose. We did not perform test-time augmentations, and the default tile overlap of 0.1 was used. The default cell probability and flow error thresholds were used, 0.0 and 0.4 respectively. For memory benchmarking, we used ‘memory profiler’ (‘mprof’ command) for CPU RAM and ‘torch.cuda.max memory allocated’ for GPU RAM - when the process used more GPU RAM than available on the GPU, the processing did slow down but was successful (e.g. the 9,600 × 9,600 image size testing).

#### Fine-tuning and evaluation

We varied the number of training images by powers of 2 (from 1 to 512 and additionally the full dataset of training images). The training images were not rescaled by the diameters of the labeled cells. The learning rate increased linearly from 0 to 1e-5 over the first 10 epochs, then decreased by factors of 2 every 10 epochs over the last 50 epochs. The network was trained for 100 epochs. Each epoch had at minimum 8 images. The batch size was set to 1. As in the whole-dataset training, the weight decay parameter was set to 0.1 and a 0.4 layer drop rate. We used the same augmentations as the previous Cellpose papers: the images were randomly rotated, flipped, and resized with a scale factor uniformly distributed between 0.75 and 1.25, and then randomly cropped to an image size of 256×256.

During test time, the 2D masks were computed as described above, without resizing the images and without test-time augmentations. For 3D mask computation, we used the 2D to 3D mask creation step described in Cellpose [3]. In brief, the flows and cell probabilities were computed on all 2D slices in XY, ZY and YZ, and then averaged to create flows in 3D, on which the dynamics steps were performed. We set the number of dynamics iterations to 1,000 and the 3D flow smoothing parameter to 2. Cell masks with fewer than 1,000 pixels were discarded.

#### Semantic segmentation training and evaluation

We retrained the Cellpose-SAM model to perform semantic segmentation on the MoNuSAC dataset, which has four classes of nuclei labeled in H&E images [24]. We added five additional output maps to Cellpose-SAM corresponding to background and the four nuclei classes, and initialized the weights of these maps with the weights from the cell probability output map, multiplied by -0.5 for the background class and 0.5 for the four nuclei classes. The loss function for the class maps was the cross-entropy loss, weighted by the inverse of the per-pixel class frequencies. The training images were not rescaled by the diameters of the labeled cells. The learning rate increased linearly from 0 to 5e-5 over the first 10 epochs, then decreased by factors of 2 every 10 epochs over the last 100 epochs. The network was trained for 500 epochs in total. The batch size was set to 16, weight decay to 0.1, and random layer drop rate to 0.4. We used the same augmentations as in the fine-tuning training.

During test time, we first computed the nucleus segmentation masks using the flows and cell probabilities as previously described. The map with the largest value across all classes was computed for each pixel, and the nucleus was assigned to the class with the most pixels within the nucleus. If this class was the background, then the nucleus mask was removed from the predicted masks. The predicted masks and classes were also shared for the challenge winners PL1, PL2, PL3 and L2 (the L1 link did not work). The winners used a hover-net architecture with a resnet, U-nets, or a feature pyramid network [69–72].

### Other models

*Cellpose cyto3* [5]: This model was trained on the Cellpose cellular dataset, the nuclear dataset, TissueNet, LiveCell, Omnipose, YeaZ, and DeepBacs, and thus we ran the model on the test set images from these datasets (excluding the YeaZ and nuclear datasets as our test splits differed from other algorithms). Two channels - cytoplasm and optionally nuclei - were used as inputs for these models. The Cellpose segmentation models are trained such that all cells and nuclei are approximately the same size in pixels across all images, by resizing each image such that the average ROI diameter is 30.0. We trained an ROI size estimation model for the cyto3 model in [5], which was used for Figure 1 and Figure S2. In Figure 3, the images were either resized as described below or the model was run directly on the image without the size model estimation. In Figure 4, as in the Cellpose 2.0 paper, the test images were rescaled using the average diameter from the training ROIs. For all analyses, the flow error threshold (quality control step) was set to 0.4, the cell probability threshold was set to 0, and test-time augmentations were on, with the tile overlap set to 0.5. In Figure 5, for all models we turned on test-time augmentations during evaluation, as in [3] and [73]. When segmenting bacterial images, we set the number of iterations ‘niter’ for the dynamics post-processing to 2,000 for all images, to improve the convergence for long and thin cells. For the fine-tuning experiments, we retrained the cyto3 model using the AdamW optimizer with a learning rate of 5e-3, weight decay of 1e-4, batch size of 8, and for 300 epochs, with the same learning rate schedule and augmentations as in the Cellpose 2.0 paper.

*CellSAM* [7]: The CellSAM model was trained on many datasets, including the Cellpose cellular dataset, TissueNet, Omnipose, DeepBacs, and MoNuSeg, on which we tested the performance of the model. We ran the ‘segment cellular image’ function available from CellSAM. CellSAM takes as input 3 channel images, with the cytoplasmic channel in blue and the nuclear channel in green. For all the test sets considered, the cells were placed in the blue channel, and the nuclear channel was optionally filled. For H&E images, the channels were averaged and input as a single channel in the blue channel as described in the paper. We used the image resizing conventions for each dataset that the authors provided in their shared datasets, available in our benchmarking script. We normalized all test images such that the minimum was set to 0 and the maximum set to 1. We enabled the histogram normalization (normalize=True) for the DeepBacs dataset as this improved performance.

*SAMCell* [9]: SAMCell trained two separate models, one on the Cellpose cellular dataset and one on the LiveCell dataset, which we applied to the Cellpose test set and the LiveCell test set respectively. We ran the ‘SlidingWindowPipeline’ ‘run’ method available from SAMCell. SAMCell only takes single channel inputs (and was trained using the grayscale version of the ‘cyto’ dataset), so we input the Cellpose test set using the average of the two channels. The Cellpose test set images were resized such that their longest side was 512, as described in their code. The LiveCell test set images were not resized and were input directly.

*MicroSAM* [8]: MicroSAM was trained on a variety of datasets, including TissueNet, LiveCell, and DeepBacs, on which we tested the performance of the model. MicroSAM was trained only on grayscale images (averaging multiple channels if available such as in TissueNet), input as the same value in all three channels. Thus, we input grayscale versions of all the test images. We used the ‘vit l lm’ model and computed the automatic instance segmentations (AIS). *PathoSAM*: PathoSAM was trained on H&E image datasets, including MoNuSeg [74], and thus we tested its performance on the MoNuSeg test set. We ran the ‘automatic segmentation wsi’ function using the ‘vit l histopathology’ model.

### Human-consensus bound

Consider annotators 1 and 2, each making mistakes relative to an absolute ground truth, or average human consensus. Consider their false positive FP_*i*_ and false negative FN_*i*_ rates relative to this consensus, and their relative rates FP_*i j*_ and FN_*i j*_ between each other, where *i, j* ∈{1, 2}, *i* ≠ *j*. It can be easily seen that FP_*i j*_ = FN _*ji*_. We assume that if the same absolute ground truth ROI is present in both the annotations of Annotator 1 and 2, then it will be matched also when comparing annotators against each other. This is not always true, but we can approximately assume it is true for the relatively low IoU threshold we require to considered two ROIs matched (0.5).

Under these conditions, FP_*i j*_ ≤FP _*j*_ + FN_*i*_, because the false positives of Annotator *j* compared to annotator *i* originate from either mistakes of Annotator *j* where a new ROI is introduced compared to ground truth (FP _*j*_) or mistakes of Annotator *i* where an ROI from the ground truth was omitted (FN_*i*_) but it was not omitted by Annotator *j* so it appears as a false positive of Annotator *j*. Furthermore, there are no other kinds of mistakes that can be counted in the FP_*i j*_. If the Annotator 1 and 2 happen to make some of the same mistakes compared to the absolute ground truth (same false positives or same false negative), then the inequality becomes strict: FP_*i j*_ *<* FP _*j*_ + FN_*i*_. Note however that the absolute ground truth is defined as a consensus across a large number of annotators, so on average the mistakes will only be identical at a rate that is the square of the single annotator mistake rates (10% error rates becomes 1% common error rates between two annotators). If we ignore this small overlap in errors, we get an approximate equality FP_*i j*_ ≈ FP _*j*_ + FN_*i*_, and since *FN*_*i j*_ = FP _*ji*_, we get

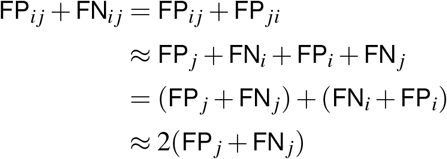

where the last approximation arises from the symmetry of considering two random annotators from the same distribution of annotators. Thus, the inter-annotator errors are approximately twice the errors between each annotator and a hypothetical “mean” or consensus annotator.

This bound may never exactly be achieved due to the approximations above, and in practice we should expect the best possible scores to be somewhere between inter-annotator performance and half of this value (where Cellpose-SAM is Figure 2c). An additional confound is that the models are trained exclusively on data from a single annotator. The ability to match segmentations from a different annotator thus is an indication of its generalization.

### Segmentation benchmarks

We tested Cellpose-SAM on the test set images from the following datasets: Cellpose, TissueNet, LiveCell, Omnipose, DeepBacs, and MoNuSeg.

We make a note about comparisons between models from different research groups. For this study, we chose to exclusively compare against models that were trained by their respective teams, and to report performance exclusively on those datasets where the models were trained with the same train/test splits (Figure 2, Figure S2). This removes the possibility of major errors when re-training a model designed by a different group. In contrast, other studies like [7] and [75] retrain the Cellpose model. To further facilitate comparisons for future studies, we will publicly release the combined dataset that was used in this study, with train/test splits for every dataset upon publication of the study. We have previously reported separately [10] on the issues of training Cellpose in [75]. Note that [7] reports better performance compared to the old Cellpose3 model (“cyto3”), but these are either reported on datasets Cellpose3 has not been trained on, or retrained by the authors from scratch. When we directly compare cyto3 to CellSAM on datasets both have been trained on, cyto3 performs better on every dataset. As described above, Cellpose-SAM performs better still, and by a substantial margin for the Cellpose test images.

#### Color and size invariance

To test color invariance we permuted the channels of images from the Cellpose test set to RGB, BRG and GBR, and also performed a random permutation per image (‘Random’), and then quantified the segmentation quality for each image (Figure 3a). To test size invariance, we resized images in the Cellpose test set such that the ROI diameters were set to 10, 15, 30, 60 and 90 (Figure 3b). We ran Cellpose-SAM, Cellpose cyto3, and CellSAM on these images without image resizing.

#### *Robustness to image degradation* as

We tested robustness to image degradation using the images from the Cellpose test set (Figure 3c-f). We added three levels of degradation for each of the four degradation type. For Poisson noise we multiplied the image by scaling factors of 5, 2.5, and 0.5 and used this as the mean for the Poisson distribution to randomly sample from for each pixel. For blurring, the Gaussian standard deviations were 2, 4, and 8, and we generated the images with Poisson noise after multiplying by 120 (small amount of Poisson noise degradation). For varying pixel size and anisotropic downsampling, the images were rescaled such that the diameter of the cells was 30 pixels for each image before downsampling. The pixel size downsampling factors were 2, 5, and 10, and the images were blurred with a Gaussian with a standard deviation of half the downsampling factor. The anisotropic downsampling factors were 2, 6, and 12 along one dimension, with blurring with a Gaussian with a standard deviation of half the downsampling factor along the same dimension. After downsampling, the images were bilinearly interpolated to their original size to be input to the networks. After these operations, each of the degraded images was normalized such that 0 was the first percentile and 1 was the 99th percentile, as in the Cellpose3 paper [5].

The noisy and blurry images were at their original size with varied sizes of cells in pixels. As in the Cellpose3 paper, for the ‘cyto3’ network and the restoration networks, we first resized these images such that the cells were 30 pixels in diameter, as we did not train size models on the degraded images. The images were not resized for Cellpose-SAM or CellSAM.

### Quantification of segmentation quality

As described in Cellpose 1 and 2, we quantified the predictions of the segmentation algorithms by matching each predicted mask to the ground-truth mask that is most similar, as defined by the intersection over union metric (IoU) between the predicted and ground-truth. We used an IoU threshold of 0.5 for all analyses in the paper. The error rate for each test image is defined using the true positives (matches with IoU above a threshold of 0.5), false positives (predicted masks without matches), and false negatives (missed ground-truth masks):

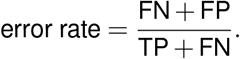

The average precision is defined for each test image

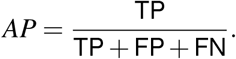

The error rates and average precisions were reported per image, with the full distribution across images shown in each violin plot.

For the semantic segmentation performance, the error rate and average precision were computed per image and per class, resulting in four scores per image, and then these four scores were averaged across classes. If an image did not contain any ground-truth masks of a certain class, then the error rate and average precision for that class were not used in the average across classes.

### Datasets

#### Main retraining

We used 18 publicly available datasets for training Cellpose-SAM. The sampling probability of each image varied depending on the image set: PhC yeast images and fluorescent bacterial images were sampled at a probability of 1% each; bright-field yeast images, phase bacterial images, and DeepBacs images at 2% each; livecell images at 5%; tissuenet images at 8%; nuclei images at 20%; and cyto2 images at 59%. We upweighted images in the cyto2 and nuclei training sets because they contained the most variability across images.

- *Cellpose (updated) dataset* : The cyto2 training dataset contains 796 training images from various sources [5, 76–80]. The dataset is available at https://www.cellpose.org/dataset. The original test dataset contained 13 images of non-biological structures, that we remove from evaluation. We instead added 12 new test images that were segmented by our annotator after the Cellpose dataset was originally published, resulting in 67 test images for the updated Cellpose test set.
- *Cellpose nuclei dataset* : This dataset of nuclear images was described in detail in [3]. It consists of 1025 training images from various sources, with about half of the images originating from the 2018 DataBowl competition [17, 18, 23, 81].
- *TissueNet* [2]: The TissueNet dataset consists of 2601 training and 1249 test images collected using fluorescent microscopy on 6 tissue types with labeled cells and nuclei (https://datasets.deepcell.org/) – we used the cellular segmentations for all images.
- *LiveCell* [1]: The LiveCell dataset consists of 3188 training and 1516 test images of 8 different cell lines collected using phase-contrast microscopy (ht tps://sartorius-research.github.io/ LIVECell/). Overlapping mask regions were removed, as described in the Cellpose 2.0 paper [4]. Many images in this dataset are incompletely annotated, which is reflected in the high error rates from Figure S2.
- *Omnipose* [19]: The Omnipose dataset consists of fluorescent bacterial images (143 training and 75 test images), and phase-contrast microscopy bacterial images (249 training and 148 test images).
- *YeaZ* [20]: The YeaZ dataset consists of bright-field and phase contrast images of yeast cells. We used 16 2D images from the phase contrast dataset for training, and 229 images from the bright-field dataset for training.
- *DeepBacs* [21]: We used the following segmentation datasets from DeepBacs: *S. aureus* bright-field and fluorescence with 56 training patches and 10 test images [82], *E. coli* bright-field with 19 training images and 15 test images [83], and *B. subtilis* fluorescence with 80 training images and 10 test images [84]; in total this is 155 training images and 35 test images.
- *Neurips 2022 challenge dataset* [22]: The Neurips 2022 challenge training dataset consists of 1,000 images with labeled cells from bright-field, fluorescent, phase-constrast and differential interference contrast imaging modalities (https://neurips22-cellseg.grand-challenge.org/neurips22-cellseg/). There are ∼10 categories of images, with each containing between 5-200 homogeneous image types. Also, a large fraction of these images are annotated sparsely. We manually selected a subset of 504 images for training, which contained more dense annotations, and with the aim of equalizing to some degree the number of images across image types. We did not evaluate performance on the validation or test set for this dataset, since the test images have the same biases and homogeneities as the training set.
- *MoNuSeg* [23]: The MoNuSeg dataset consists of 37 training images and 14 test images with labeled nuclei, available at https://monuseg.grand-challenge.org/. 30 of the training images were also included in our Cellpose Nuclei train/test datasets in grayscale and inverted - we removed them for the visualization in Figure 2a.
- *MoNuSAC* [24]: The MoNuSAC dataset consists of H&E images from 4 different organs with labeled nuclei and nuclei classification, with 209 training images and 85 test images, available at https://monusac-2020.grand-challenge.org/. The main model was only trained on the training images. The nuclei classification labels from the training images were used to train the semantic segmentation model described in Figure 5, with the performance on test images shown in the figure.
- *CryoNuSeg* [25]: The CryoNuSeg dataset consists of H&E images from 10 different human organs with labeled nuclei (https://www.kaggle.com/datasets/ipateam/segmentation-of-nuclei-in-cryosectioned-he-images). We used all available images (30) as training images.
- *NuInsSeg* [26]: The NuInsSeg dataset consists of H&E images from 31 different human and mouse organs with labeled nuclei, available at https://www.kaggle.com/datasets/ipateam/nuinsseg. We used all available images (665) as training images.
- *BCCD* [27]: The blood cell segmentation dataset consists of blood smear images taken with a light microscope (https://www.kaggle.com/datasets/jeetblahiri/bccd-dataset-with-mask). There are 1,169 training images in the dataset, all of which were used for training.
- *CPM 15+17 and TNBC* [28, 29]: CPM 15 and 17 and TNBC consist of H&E images with labeled nuclei. CPM 15 + 17 are from brain cancer patients, with 15 and 32 training images per dataset respectively. TNBC consists of 50 images from triple negative breast cancer patients, all of which were used for training. The datasets are available at https://drive.google.com/drive/folders/1l55cv3DuY-f7-JotDN7N5nbNnjbLWchK.
- *LynSec* [30, 31]: LynSec consists of 699 IHC and H&E images from lymphoma patients with labeled nuclei (https://zenodo.org/records/8065174). We chose 616 of these images randomly to use in our training set.
- *IHC TMA* [32, 33]: The IHC TMA dataset consists of TMA sections from non-small cell lung cancer patients with labeled nuclei (https://doi.org/10.5281/zenodo.7647846). There are 195 and 36 images in the training and validation sets, we used all of these for training.
- *CoNIC* [34]: The CoNIC dataset consists of 4,981 H&E images with labeled nuclei and nuclei classification (https://www.kaggle.com/datasets/aadimator/conic-challenge-dataset?select=data). We randomly chose 3,863 of the images for training, all of which had at least one labeled nuclei (a subset of images in the dataset had no labels). We used all nuclei labels.
- *PanNuke* [35, 36]: The PanNuke dataset consists of 7,898 H&E images from 19 tissues types from cancer patients with labeled nuclei and nuclei classification (https://warwick.ac.uk/fac/cross_fac/tia/data/pannuke). We randomly chose 6,053 of the images for training, all of which had at least one labeled nuclei (a subset of images in the dataset had no labels). We used all nuclei labels.

For testing we only used the datasets with a well-defined test set, as described above, and on which other models had been trained, because those were the only datasets in which we could ensure train/test splits were consistent across available models.

#### Fine-tuning datasets

We tested fine-tuning performance on three publicly available datasets:

- The lateral root dataset consists of 27 volumes from three different *Arabidopsis thaliana* lateral root primordia timelapse recordings, acquired every 30 minutes with a voxel size of 0.1625 × 0.1625 × 0.250 um (X x Y x Z), with ground-truth 3D segmentations
- (https://osf.io/2rszy/). We reduced the size of the volumes by a factor of 4 in XY and by a factor of 2.6 in Z (to create an isotropic volume). We used all 17 of the volumes from Movies 1 and 3 for training, and two of the ground-truth volumes from Movie 2 for testing, which were in the original test set (timepoint 10 and 20). We divided by movie id to avoid any relationship between the train and test set. From each training and test stack we took 40 slices, 20 in XY and 10 in each ZY and ZX. These slices were evenly spaced within the volume, with a padding of 10 on each side of the dimension being sampled. This resulted in 680 2D training slices and 80 2D test slices. We tested 3D segmentation using the 3D ground-truth for the two 3D test volumes.
- The ovule dataset consists of 31 volumes from *Arabidopsis thaliana* ovule recordings with a voxel size of 0.075 × 0.075 × 0.235 um, with ground-truth 3D segmentations [44] (https://osf.io/w38uf/). We reduced the size of the volumes by 1.33 in XY and by a factor of 2.35 in Z (to create an isotropic volume). We used the 22 defined training volumes for the training set and the 7 defined test set volumes for testing. As in the lateral root dataset, we took 40 2D slices from each stack, resulting in 880 2D training slices and 280 2D test slices, and tested 3D segmentation on the 7 full volumes.
- The BlastoSPIM dataset consists of 231 3D volumes of early-stage mouse embryos from 55 different embryos, and 80 3D volumes of late-stage mouse embryos (‘Blast’) acquired on a confocal microscope, with a voxel size of 0.208 × 0.208 × 2.0 um [43]. We defined the test set as the volumes from 5 random early-stage embryos (16 volumes in total), and 10 random volumes from the late-stage embryo dataset. The training set contained volumes from the other 50 early-stage embryos and 70 Blast volumes. We upsampled the volumes in Z by a factor of 10 to create isotropic volumes. We used 3 2D slices from each training and test volume, 1 in XY, ZY and ZX. These slices were sampled randomly, until a slice with at least one mask was obtained, with a padding of 30 pixels on each side of the dimension being sampled. This resulted in 855 2D training slices and 78 2D test slices. We tested 3D segmentation on the 26 full 3D test volumes.

## Footnotes

1 https://github.com/MouseLand/cellpose

2 https://huggingface.co/spaces/mouseland/cellpose

---

**Table S1:** GPU statistics for inference. Runtimes and memory profiling for model inference on a consumer GPU (RTX 4070S, $600, 12 GB of GPU RAM) on Windows with a maximum batch size of 32 for different images sizes.

| image size (pixels) | file size | GPU RAM used (reported) | RAM used (reported) | batch size | runtime |
| --- | --- | --- | --- | --- | --- |
| 150 | 270KB | 2.45 GB | 2.18 GB | 1 | 0.37 sec |
| 300 | 1.08MB | 2.45 GB | 2.18 GB | 4 | 0.41 sec |
| 600 | 4.32MB | 3.39 GB | 2.23 GB | 9 | 0.87 sec |
| 1,200 | 17.2MB | 8.90 GB | 3.57 GB | 32 | 3.24 sec |
| 2,400 | 69MB | 8.90 GB | 3.84 GB | 32 | 12.48 sec |
| 4,800 | 276MB | 8.90 GB | 4.80 GB | 32 | 46.71 sec |
| 9,600 | 1.11GB | 12.28 GB | 18.40 GB | 32 | 368.88 sec |

**Table S2:** GPU statistics for training. Runtimes and memory profiling for model training with a small number (8) of user provided annotated images with a batch size of 1, epoch size of 8, and 100 epochs. Typical for the human-in-the-loop retraining stage.

| GPUs | MSRP | GPU RAM max | GPU RAM used (reported) | RAM used (reported) | runtime |
| --- | --- | --- | --- | --- | --- |
| RTX 4060 | \$300 | 8GB | 8.53 GB | 3.01 GB | 363.58 sec |
| T4 (colab) | free | 16GB | 8.53 GB | 1.81 GB | 185.17 sec |
| RTX 4070S | \$600 | 12GB | 8.53 GB | 1.87 GB | 55.08 sec |
| RTX 5090 | \$2,000 | 32GB | 8.53 GB | 2.56 GB | 33.71 sec |
| A100 | >\$10,000 | 80GB | 8.53 GB | 2.26 GB | 27.30 sec |

**Figure S1:**
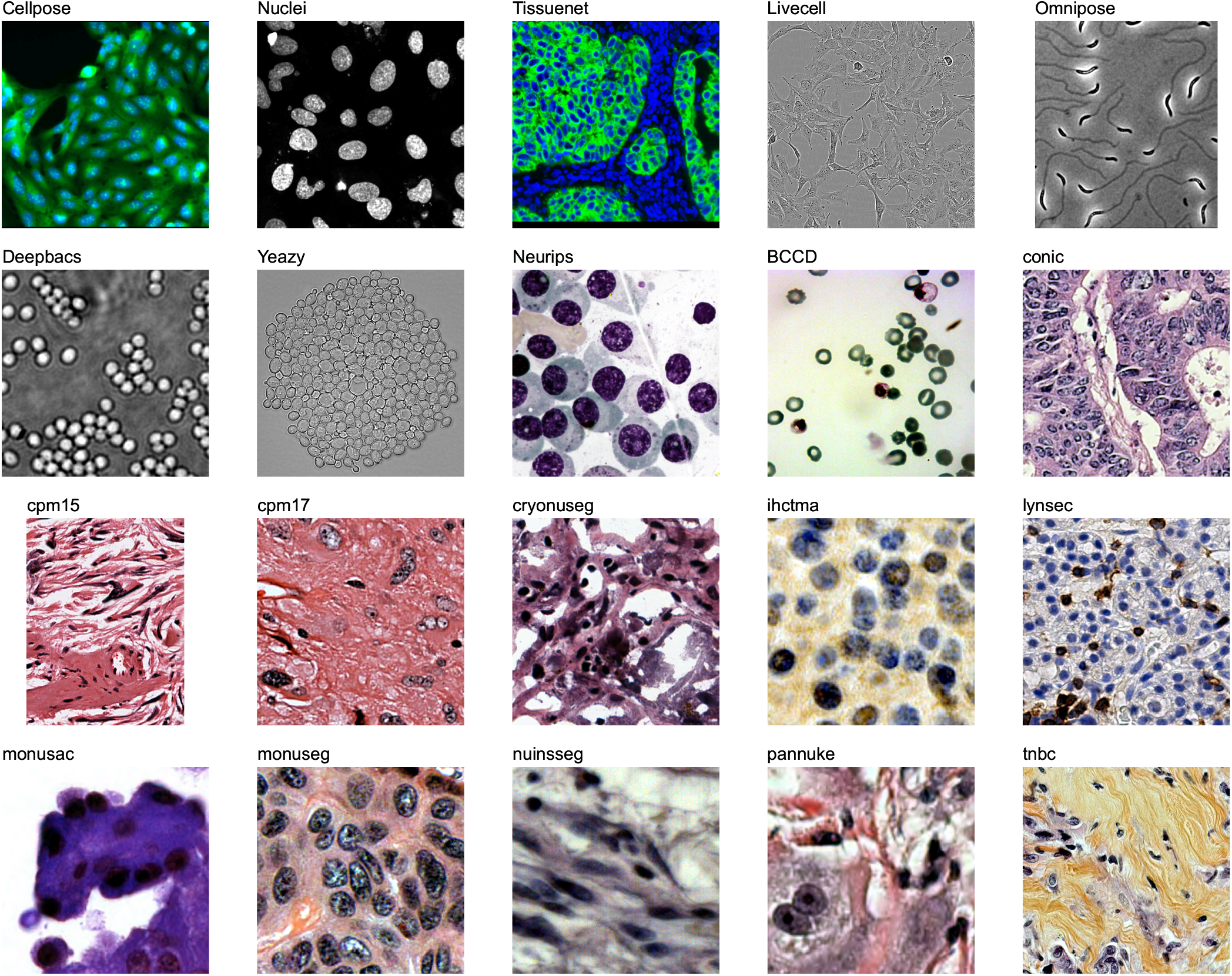
Example training images. Images originate from a total of 20 distinct datasets. For the Cellpose dataset, the individual image is not representative of the heterogeneity of the dataset (see Figure 2a).

**Figure S2:**
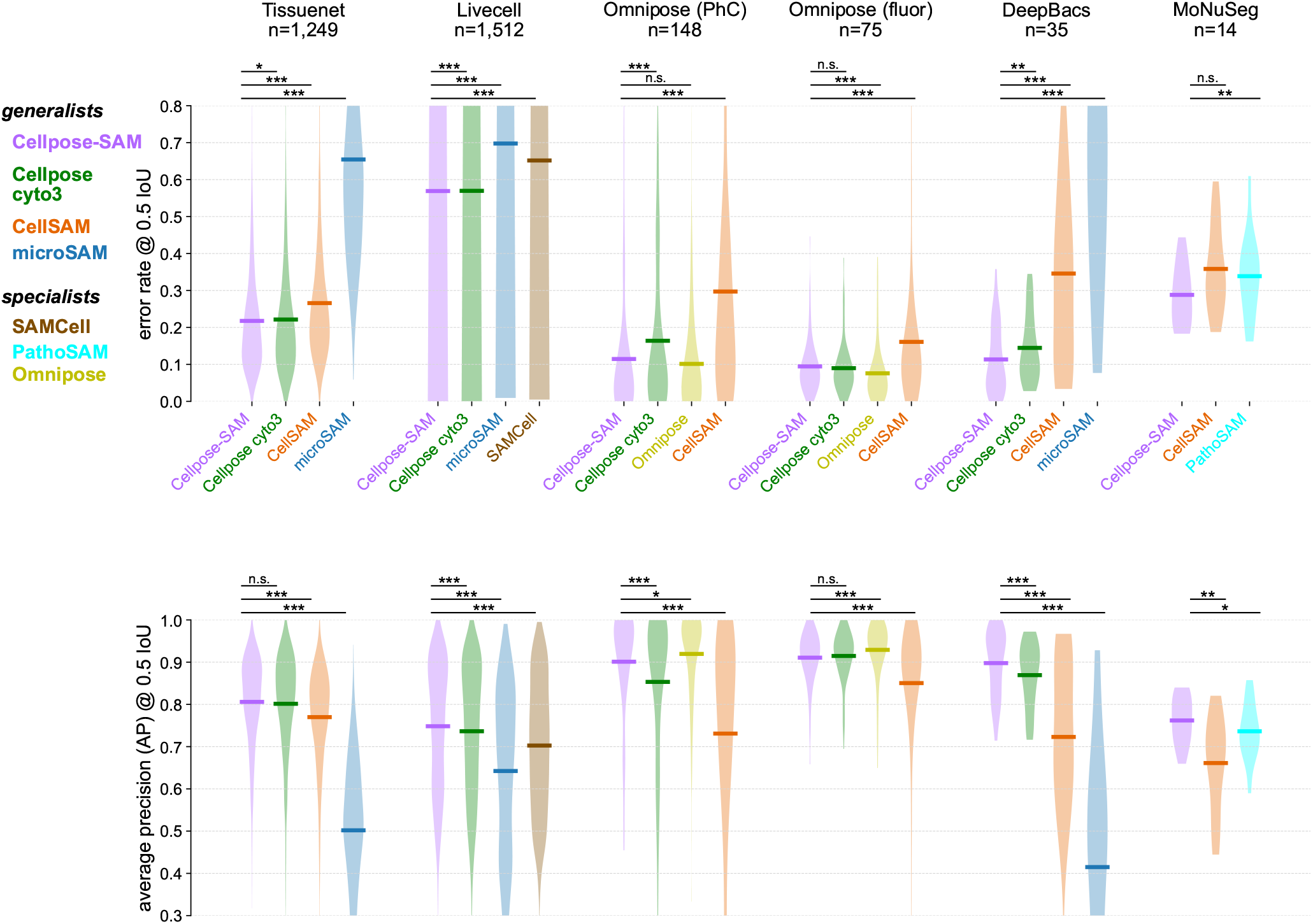
Segmentation performance on other datasets. Similar to Figure 2de. Performance is reported with either error rates (top) or AP @ IoU = 0.5 (bottom). Algorithms are included in each panel if : 1) they have been trained on the respective dataset; 2) they used the publicly available train/test split. We did not retrain any of the algorithms from other groups. Specialist models were only trained on a single dataset. Wilcoxon signed-rank test performed - CellposeSAM performs as well as (n.s.) or significantly better than all other algorithms, besides the specialist Omnipose model on the Omnipose (fluor) dataset.

